# Bacterial but not fungal rhizosphere communities differ among perennial grass ecotypes under abiotic environmental stress

**DOI:** 10.1101/2021.11.15.468685

**Authors:** Soumyadev Sarkar, Abigail Kamke, Kaitlyn Ward, QingHong Ran, Brandi Feehan, Shiva Thapa, Lauren Anderson, Matthew Galliart, Ari Jumpponen, Loretta Johnson, Sonny T M Lee

## Abstract

Environmental change, especially frequent droughts, is predicted to detrimentally impact the North American perennial grasslands. Consistent dry spells will affect plant communities as well as their associated rhizobiomes, possibly altering the plant host performance under environmental stress. Therefore, there is a need to understand the impact of drought on the rhizobiome, and how the rhizobiome may modulate host performance and ameliorate its response to drought stress. In this study, we analyzed bacterial and fungal communities in the rhizospheres of three ecotypes (dry, mesic, and wet) of a dominant prairie grass, *Andropogon gerardii*. The ecotypes were established in 2010 in a common garden design and grown for a decade under persistent dry conditions at the arid margin of the species’ range in Colby Kansas. The experiment aimed to answer whether and to what extent do the different ecotypes maintain or recruit distinct rhizobiomes after ten years in an arid climate. In order to answer this question, we screened the bacterial and fungal rhizobiome profiles of the ecotypes under the arid conditions of western KS as a surrogate for future climate environmental stress using 16S rRNA and ITS2 metabarcoding sequencing. Under these conditions, bacterial communities differed compositionally among the *A. gerardii* ecotypes, whereas the fungal communities did not. The ecotypes were instrumental in driving the differences among bacterial rhizobiomes, as the ecotypes maintained distinct bacterial rhizobiomes even after ten years at the edge of the host species range. This study will aid us to optimize plant productivity through the use of different ecotypes under future abiotic environmental stress, especially drought.

## Introduction

The rhizosphere, which plays a pivotal role in plant function by facilitating elemental and water cycling, and uptake of nutrients (Adl 2016), is densely populated by diverse microbial communities – the rhizobiome. A wide range of complex interactions ranging from symbiotic to competitive among the rhizobiome microorganisms govern the carbon, nitrogen, and phosphorus uptake and transformations (Ahkami et al. 2017; Barea et al. 2005). These interactions exist not only among the microorganisms but also between the plant hosts and their associated rhizobiome. For example, plant hosts may selectively attract and/or repel specific soil microbial communities through their root exudates (Bais et al. 2001; Estabrook and Yoder 1998). Furthermore, these microorganisms may establish symbiotic relationships with the host plants, safeguarding the host against pathogens (Nardi et al. 2000; Jones et al. 2019). These interactions between the plant host and its rhizobiome are likely to be highly specific and ultimately important for community stability, ecosystem functioning, and maintaining soil biodiversity (Wagg et al. 2021).

Global change can have adverse effects on microbe-microbe and plant-microbe interactions, which in turn, can impact the ecology of the rhizosphere and ecosystem function (Ahkami et al. 2017). Some studies have focused on the impacts of climate change on rhizospheres (Classen et al. 2015; Bei et al. 2019; Compant, Van Der Heijden, and Sessitsch 2010). However, more concerted efforts are needed to fill the knowledge gaps between how rhizobiomes may be influenced by the interactive effects of the plant host and the ever-changing climate. Ultimately, this will help us to maximize plant growth and survival in stressful environments (Long and Zhu 2009; Long, Marshall-Colon, and Zhu 2015; Ort et al. 2015), especially impacts from drought.

Microorganisms in the rhizosphere are sensitive to environmental conditions (e.g., Rudgers et al. 2021) and can be good indicators of soil quality (Mendes, Garbeva, and Raaijmakers 2013). Dissecting the rhizosphere bacterial and fungal communities, their functional roles, and their interactions with the plant-hosts are crucial to developing future methods to improve drought tolerance and to improve plant productivity. Climate change, characterized by rising temperature and shifted precipitation patterns, has caused the increase in drought frequency (Wanders and Wada 2015; Huang et al. 2017) and severity (Liu et al. 2015; Ayantobo et al. 2017). Thus, comprehending the extent of the variability among ecotypes of the dominant species and how this variability interacts under the predicted arid conditions is important. This is especially critical because the dominant species greatly impact ecosystem processes such as carbon assimilation, nutrient cycling etc. (Grime 1998). The motivation to understand plant host-microbe interaction in dominant grasses is even more crucial in the case of tallgrass prairies, in which grasses are responsible for the majority of the carbon fixing, nutrient cycling and biomass (Risser, Birney, and Blocker 1981; Johnson and Matchett 2001).

Our studies focus on Big Bluestem, *Andropogon gerardii*, the dominant native grass species in the tallgrass prairies of central North America (Gustafson, Gibson, and Nickrent 2004). *Andropogon gerardii* is widely distributed across the Midwest and Northeastern US (USDA database). Our study focuses on the Central grasslands where this grass dominates, stretching from western Kansas to southern Illinois (Johnson et al. 2015). This precipitation gradient includes a semiarid environment, a region of intermediate precipitation, and a region of heavy rainfall (Olsen et al. 2013). Galliart and colleagues have demonstrated that within this steep precipitation gradient, three genetically distinct *A. gerardii* regional climate ecotypes (dry, mesic, and wet) exist (Gray et al. 2014; Galliart et al. 2019), in terms of leaf area, height, and blade width (Galliart et al. 2020), allocation to roots (Mendola et al. 2016), and chlorophyll abundance (Caudle et al. 2014). Climate change models have also predicted a strong phenotypic cline in *A. gerardii* across this longitudinal precipitation gradient (Galliart et al. 2019, 2020; Smith et al. 2017). However, we have limited understanding on how the three ecotypes and their rhizobiomes are differently affected by a semiarid environment to which some ecotypes may be more adapted to than others. As such, understanding the shifts and interaction between the *A. gerardii* plant host and its associated microbiome at Colby Kansas will provide insights into exploring the impact of future climate change on grasslands and the microbiome.

The goal of this study was to analyze the composition of bacterial and fungal communities in the rhizobiomes of the dry, mesic and wet *A. gerardii* ecotypes originating in Hays Kansas (rainfall ~500 mm/year), Manhattan Kansas (rainfall ~870 mm/year) and Carbondale Illinois (rainfall ~1,200 mm/year), respectively. All ecotypes were planted in Colby Kansas (rainfall ~500 mm/year) and grown for ten years prior to sampling. We asked to what extent do ecotypes of a dominant prairie grass maintain or recruit distinct rhizobiomes after ten years of growing in a semi-arid climate where precipitation is lower than where the ecotypes originated. We were specifically interested in deciphering the extent of ecotypic variation and/or pressure of a semiarid environment on the dominant tall-grass prairie grass rhizobiome. We postulated that the plant host would exert their ecotypic influences on the rhizobiome even under environmental stress, and thus would observe differences in the ecotypic microbial community. This study aims to provide insights necessary to preserve the prairie ecosystems under climate change pressures.

## Materials and Methods

### Experimental design and sampling

The common garden in Colby is located at the KSU Agricultural Research Center in Thomas County (39°23’N, 101°04’W). The common garden was established in 2010 - ten years before our sampling. The seeds of four populations of each of native dry (Hays), mesic (Manhattan KS) and wet (Carbondale Illinois) *A. gerardii* ecotypes (Galliart et al. 2020) were germinated and then grown inside the greenhouse in potting mix substrate (Metro-Mix 510). Established 3-4-month-old plants from all the populations were then planted in western Kansas (Colby) as a surrogate for the drier conditions expected in the future (Intergovernmental Panel on Climate Change 2014). Each ecotype was represented by four populations with twelve replicate plants (Galliart et al. 2020). There were a total of 12 plants (4 populations X 3 ecotypes) in a randomly complete block design with 10 blocks (Galliart et al. 2020). Plants were planted 50 cm apart along each row, and the soil around the plants was covered with a water-penetrable cloth to control unwanted plants. Some plants did not survive through the ten years in the common garden. We collected a total of 95 rhizosphere soil cores (15 cm deep, 1.25 cm diameter) from the dry (n=33), mesic (n=30) and wet (n=32) ecotypes during the growing season in Summer 2019. Each soil core was placed in a plastic bag, transported on ice and stored at −80°C until genomic DNA extraction.

### DNA extraction, metabarcoding and analyses

We extracted the genomic DNA from the root samples and soil associated with it (0.150g each) using the Omega E.Z.N.A. Soil DNA Kit (Omega Bio-Tek, Inc., Norcross, GA, USA), with a slightly modified protocol. Bulk soil was separated from the rhizosphere soil by hand shaking the roots gently, and any soil that was attached to the root was considered part of the rhizosphere. Briefly, we modified the protocol using a QIAGEN TissueLyser II (Qiagen, Hilden, Germany) for 2 minutes at 20 rev/s, and eluted the purified DNA with a final volume of 100 μl. The extracted microbial DNA were sequenced on the Illumina MiSeq, with the 16S rRNA V4 region amplified using the primers 515F and 806R with barcodes (Caporaso et al. 2012), and the internal transcribed spacer region ITS2 amplified using the primers fITS7 (Ferrer et al. 2001; Kumeda and Asao 1996; Ihrmark et al. 2012) and ITS4 (White et al. 1990), at the Kansas State University Integrated Genomics Facility.

We acquired a total of 1,729,418 bacterial and 3,570,536 fungal sequences before quality check (QC) and trimming (Supplementary Table S1). We used QIIME 2 (v. 2019.7) to process the sequence data and to profile the rhizobiome communities (Bolyen et al. 2019). We used QIIME 2 plugin cutadapt (Martin 2011) to remove the primer sequences; reads with no primer were discarded. Additionally we used DADA2 (Callahan et al. 2016) for quality control with the same parameters across different runs, and truncated the reads to length where the 25th percentile of the reads had a quality score below 15. The pre-trained classifier offered by QIIME 2, using Silva database (v. 132) was used for taxonomic assignment for bacteria. Similarly, the UNITE classifier was trained on the full reference “develop” sequences (version 8.2, release date 2020-2-20) (Community 2019) using QIIME 2 2020.2 before taxonomic assignment of the fungal reads. α-diversity was calculated to present the species diversity in each sample (Consortium and The Human Microbiome Project Consortium 2012). We estimated observed richness (S_Obs_), Shannon’s diversity (H’), and Faith’s PD using a rarefield dataset (8,023 reads for 16S; 10,258 reads for ITS). Similar to the α-diversity analysis, we used UniFrac and Bray Curtis distances to compare the community dissimilarity among the different ecotypes, and used Non-metric multidimensional scaling (NMDS) to visualize the distance matrices.

Differences in the relative abundances of bacterial and fungal phyla among the ecotypes were analyzed using ANOVA in R followed by Tukey’s post-hoc test (p < 0.05) (Team 2015). SIMPER post-hoc analyses were performed to identify those community members that contributed to the highest differences among the ecotypes. The cut-off used for all SIMPER analyses was 70%. We used DeSEQ2 and highlighted marked differences in the disproportionate relative abundance of bacteria taxa among the ecotypes (p < 0.05) (Love, Huber, and Anders 2014). All raw sequence data is available in the NCBI under the bioproject PRJNA 772708 and biosamples (SAMN 22405120 - SAMN 22405309).

## Results and Discussion

We dissected the rhizosphere bacterial and fungal communities associated with the dominant tallgrass prairie species, *A. gerardii*. We recovered an average of 17,181 + 4,935 counts per sample for bacteria, and 37,015 + 7,394 counts for fungi after primer trimming (Supplementary Table S1). Of the recovered counts, an average of 71.45% bacterial counts and 61.55% fungal counts were annotated to the species level on SILVA and UNITE respectively. Any unknown or unclassified amplicon sequence variants (ASVs) were removed from downstream analyses. The bacterial and fungal taxon assignments along with their counts in each sample are provided in Supplementary Table S2.

### No differences in bacterial or fungal α-diversity among host ecotypes

We found no support for differences in bacterial α-diversity (S_Obs_, Shannon’s H’index: H = 6.374, p = 0.041, or Faith’s PD index: H = 3.626, p = 0.163 and observed OTUs index: H = 3.959, p = 0.138) among *A. gerardii* ecotypes (Figure 1A). Venn diagrams of the shared bacterial and archaeal ASVs reveal 1,703 shared ASVs (98.66%) among the three ecotypic rhizobiomes (Figure 1A). These shared ASVs were Actinobacteria, Proteobacteria, Acidobacteria, Verrucomicrobia, Bacteroidetes, Thaumarchaeota, Chloroflexi, Firmicutes, Patescibacteria, Planctomycetes, Armatimonadetes, Gemmatimonadetes, Latescibacteria, Cyanobacteria, Rokubacteria, Entotheonellaeota, Nitrospirae, BRC1, Chlamydiae, Dependentiae, FBP, Elusimicrobia, Deinococcus-Thermus, Fibrobacteres, and WS2. Similar to the bacterial α-diversity, we observed no support for differences in the fungal rhizobiome α-diversity among the three *A. gerardii* ecotypes (S_Obs_, Shannon’s H’index: H = 3.759, p = 0.153, or Faith’s PD index: H = 4.798, p = 0.091 and observed OTUs index: H = 3.393, p = 0.183) (Figure 1B). There was one unique fungal ASV belonging to the wet ecotypes, and none in the other ecotypes. There were 829 (99.28%) overlapping ASVs among the three ecotypes (Figure 1B). The rhizobiome ASVs that were shared among the dry, wet and mesic ecotypes belonged to Basidiomycota, Ascomycota, Mortierellomycota, Glomeromycota, Kickxellomycota, Chrytridiomycota, Rozellomycota, Aphelidiomycota, and Entomophthoromycota.

**Figure 1A:**
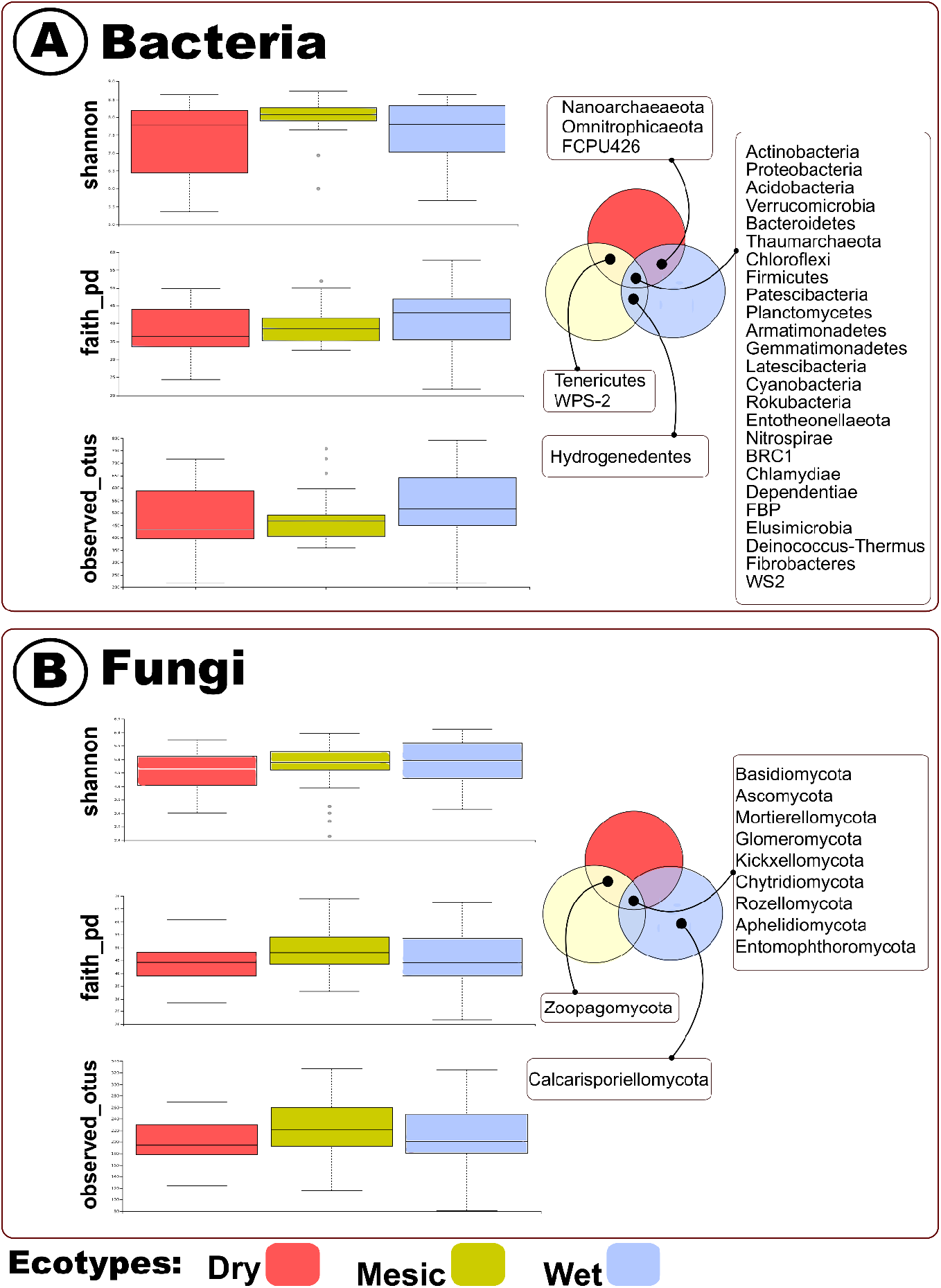
Bacterial α-*Diversity* indices among the dry, mesic, and wet ecotypes. (a) Shannon index (b) faith-pd index and (c) observed OTUs index (d) Venn diagrams representing the overlapping bacterial and archaeal ASVs among the dry, mesic, and wet rhizobiomes. The bacterial α-*Diversity* was not significantly different among the samples. Figure 1B: Fungal α-*Diversity* indices among the dry, mesic, and wet ecotypes. (a) Shannon index (b) faith-pd index and (c) observed OTUs index (d) Venn diagrams representing the overlapping fungal ASVs among the dry, mesic, and wet rhizobiomes. The fungal α-*Diversity* was not significantly different among the samples.

### Bacterial composition in the rhizobiome differs among host ecotype ecotypic rhizobiome

Bacterial communities at the phylum level differed among the three *A. gerardii* ecotypes (PERMANOVA, Pseudo-F 4.1963, p (perm): 0.001, p (MC): 0.002, NMDS; stress: 0.13). We analyzed the samples based on Bray-Curtis similarity, and observed that the mesic data cloud dispersion was smaller than that of the dry and wet ecotypes (Figure 2A). In contrast to bacteria, we did not observe any evidence for differences in the fungal rhizobiome composition among the dry, wet and mesic ecotypic rhizobiomes (PERMANOVA, Pseudo-F 2.0827, p(perm): 0.071, p(MC): 0.08, NMDS; stress 0.11; Figure 2B).

**Figure 2:**
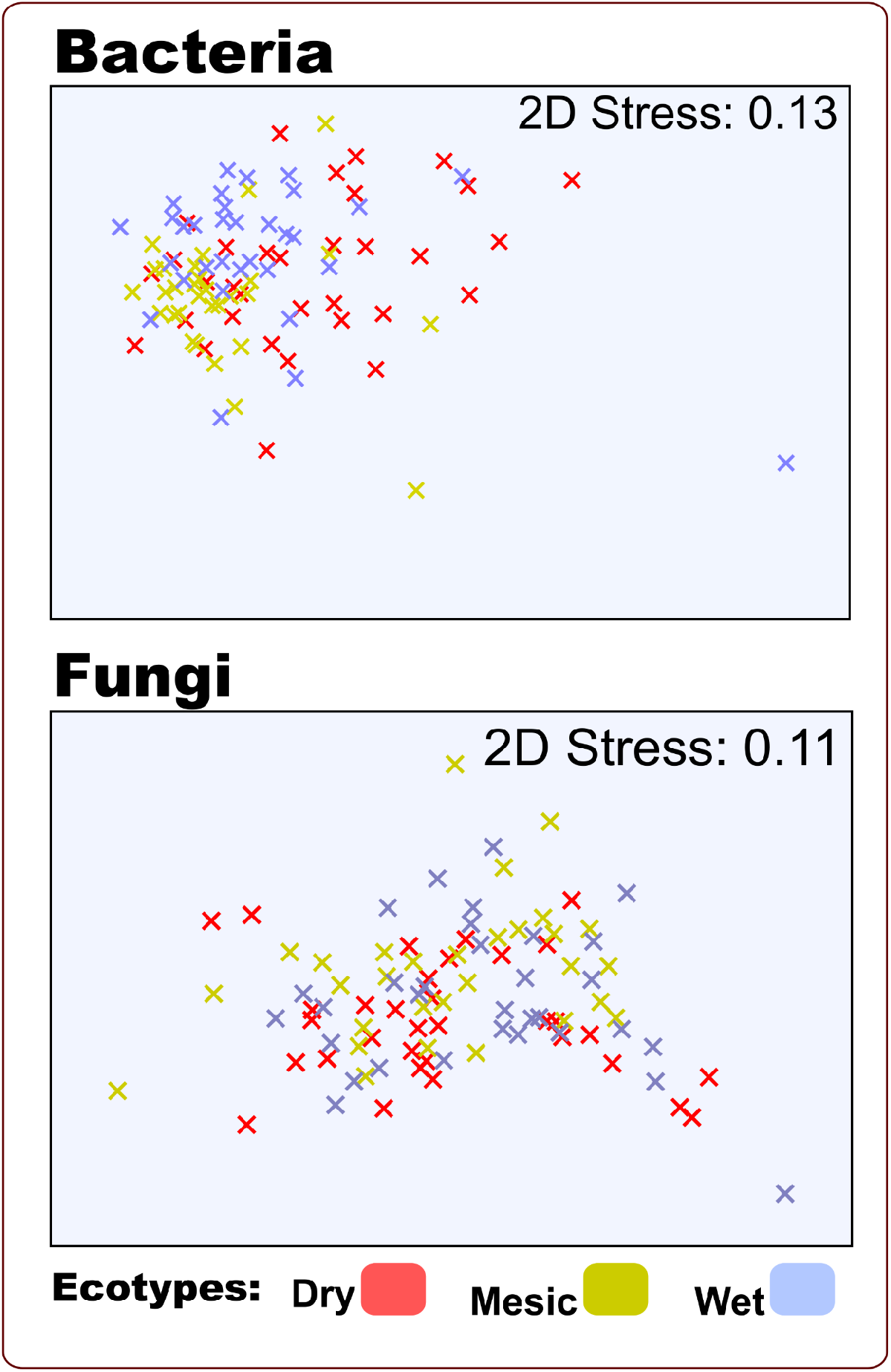
Bacterial and fungal *β-Diversity* between the dry, mesic, and wet ecotypes. NMDS ordinations were obtained from Bray-Curtis similarity matrix. The matrix was calculated from square-root transformed relative abundance of 16S and ITS rRNA sequences (A) bacterial community compositions are separated between the three ecotypes (B) fungal community compositions are not separated between the three ecotypes.

### Significant differences in ecotypic bacterial diversity and structure

Based on pairwise Kruskal-Wallis test, bacterial rhizobiome communities were distinct among all three ecotypes (dry v wet: p (MC): 0.034, dry v mesic: p (MC): 0.001, wet v mesic: p (MC): 0.005). The top seven bacterial phyla present in all the three ecotypes are Proteobacteria, Actinobacteria, Acidobacteria, Chloroflexi, Bacteroidetes, Verrucomicrobia, Planctomycetes, and one archaeal taxa, Thaumarchaeota (Figure 3, Supplementary Figure S1). Acidobacteria, Bacteroidetes, and Proteobacteria are ubiquitous in soil, suggesting their importance in our samples (Chowdhury et al. 2019; Sarkar et al. 2020). Similarly, Chloroflexi, Verrucomicrobia, Planctomycetes, and Thaumarchaeota are common soil dwellers, reported to contribute to diverse soil processes (Zhang, Chen, and Han 2013; Navarrete et al. 2015; Ivanova et al. 2016; Lu, Seuradge, and Neufeld 2017), which also corroborates with the detection of the bacteria in our analyses regardless of the ecotypes. We used ANOVA followed by Tukey post-hoc test (Team 2015), and showed that the relative abundance of Proteobacteria (F=7.292, p: 0.004) and Thaumarchaeota (F=4.451, p: 0.020) differed between the dry and wet ecotypes. Proteobacteria were more abundant in the dry than in the wet ecotype, whereas Thaumarchaeota abundance was the opposite (Figure 3). Some Proteobacteria may improve plant performance and growth, and can increase in abundance under drought conditions (Jang et al. 2020; Kim et al. 2011), suggesting that Proteobacteria might be important for the sustainable growth of *A.gerardii* under the challenging environmental conditions in Colby. Thaumarchaeota are the dominant archaea in soil systems (Schleper and Nicol 2010), and well-known ammonia oxidizers (Stieglmeier, Alves, and Schleper 2014). We surmise that Thaumarchaeota in our study might have the potential to enhance the resilience of the *A.gerardii* wet ecotype under abiotic stressful conditions through the transformation of ammonia into nitrate (Taffner et al. 2018).

**Figure 3:**
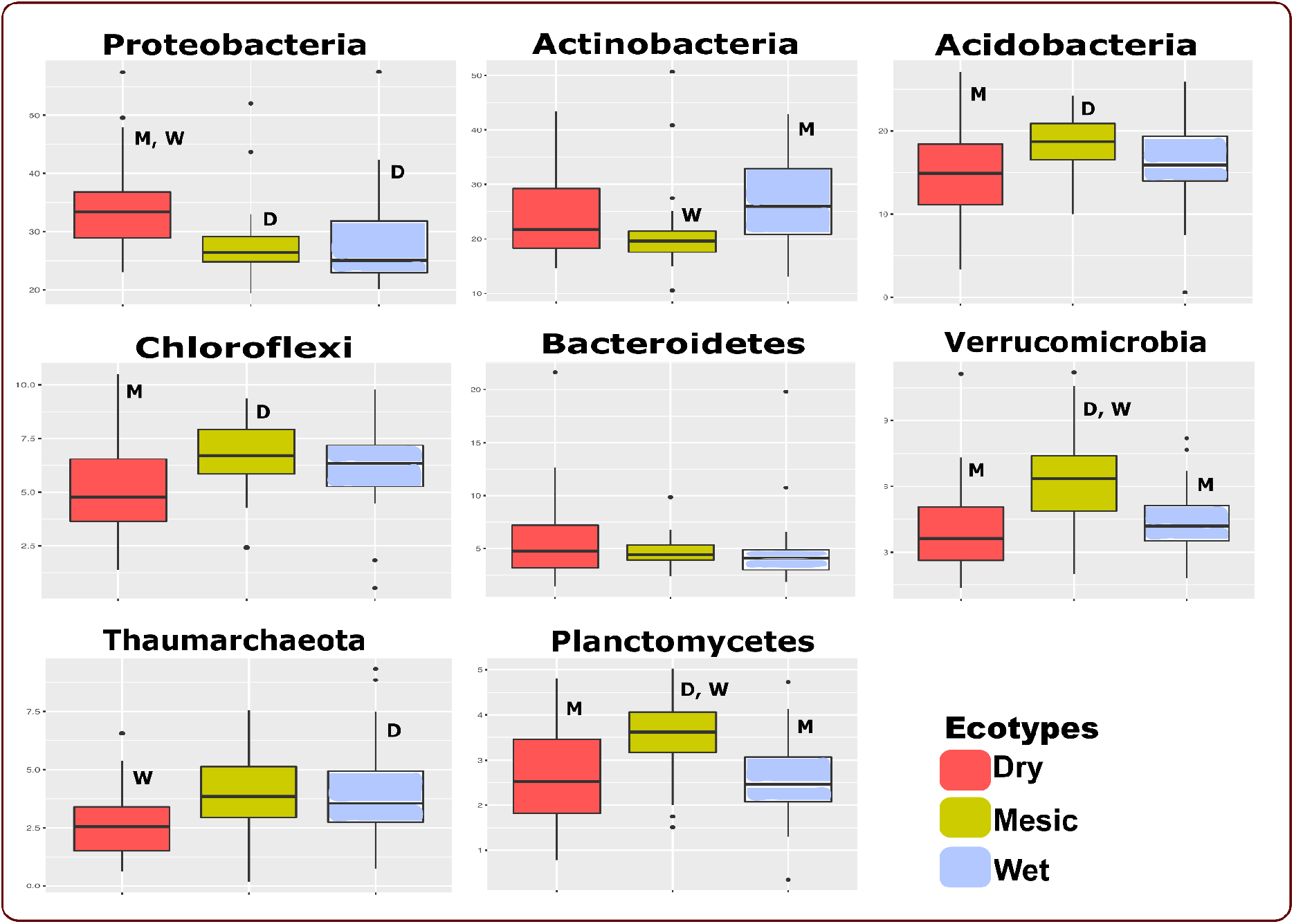
The relative abundance of the top seven bacterial and one archaeal taxa present in all the three ecotypes. Proteobacteria and Thaumarchaeota were significantly different between the dry and wet ecotypes. Letters in a box plot are significantly different at p < .05 (D, M, W - significantly different from dry, mesic and ecotypes respectively).

Post-hoc SIMPER analyses at the Phylum level showed that Proteobacteria, Actinobacteria, Acidobacteria, Chloroflexi, Bacteroidetes and Verrucomicrobia contributed most to the differences among the dry, mesic and wet ecotypes (Supplementary Table S3). Using post-hoc SIMPER analyses, we observed that Actinobacteria, Acidobacteria, and Proteobacteria contributed to the greatest differences between the dry and mesic ecotypes, dry and wet ecotypes, and mesic and wet ecotypes. Verrucomicrobia and Bacteroidetes contributed to the greatest differences between the dry and mesic ecotypes (Supplementary Table S3). Planctomycetes and Thaumarchaeota contributed to the differences only in the mesic and wet ecotypes respectively (Supplementary Table S3). Comparing the dry and wet ecotypes, we observed that Bacteroidetes, and Thaumarchaeota contributed to their greatest differences. Verrucomicrobia, and Thaumarchaeota contributed most to the differences between the mesic and wet ecotypes.

We analysed the data with DeSEQ2 (p < 0.05, Figure 4) and detected that *Rhizobium* had significant differences between the dry and wet ecotypic rhizobiomes. *Rhizobium* had higher relative abundances in the dry ecotype. Comparing dry and mesic ecotypes, we noticed that *Rhizobium, Pseudomonas, Cellulomonas, Rhodococcus, Parviterribacter, Parasegetibacter*, *Flavihumibacter, Cellvibrio*, and *Candidatus Berkiella* were more dominant in dry ecotype than in mesic ecotype. On the opposite side, *Microbispora, Sorangium, Zavarzinella,* and *Candidatus Udaeobacter* had more dominance in mesic than in wet ecotype. In other studies, *Rhizobium* has been found to be drought-stress tolerant (Rehman and Nautiyal 2002), and well-known to aid plants during drought conditions (Staudinger et al. 2016). Putting it all together, our study suggested that *Rhizobium*, being the most predominant in the dry ecotype, might have the potential influence to benefit the host in the dry environments. This may also help to explain the higher leaf nitrogen concentrations and higher chlorophyll absorbance we observed in the dry ecotype, regardless of planting location (Caudle et al.2014, Galliart et al. 2020). We compared the differences between mesic and wet ecotypes as well, and observed that *Parafrigoribacterium, Ktedonobacter, Streptosporangium, Acidicapsa*, and *Pseudoduganella* were more dominant in the mesic ecotype. On the contrary, the top genera that were more predominant in wet ecotypes as compared to the mesic belong to *Candidatus Berkiella, Cellvibrio, Flavihumibacter, Terrabacter, Parasegetibacter, Parviterribacter, Cellulomonas, Solitalea, Sphaerisporangium, Nonomuraea, Achromobacter, Acinetobacter, Pseudorhodoferax, Nocardia, Leucobacter*, among others (Supplementary Table S4). *Leucobacter* has been identified to grow in wet, low-light environments (Muir and Tan 2007), and we observed that *Leucobacter* had higher relative abundance in the wet ecotype (Muir Rachel E. and Tan Man-Wah 2008) suggesting that *Leucobacter* might be better adapted to the wet environment.

**Figure 4:**
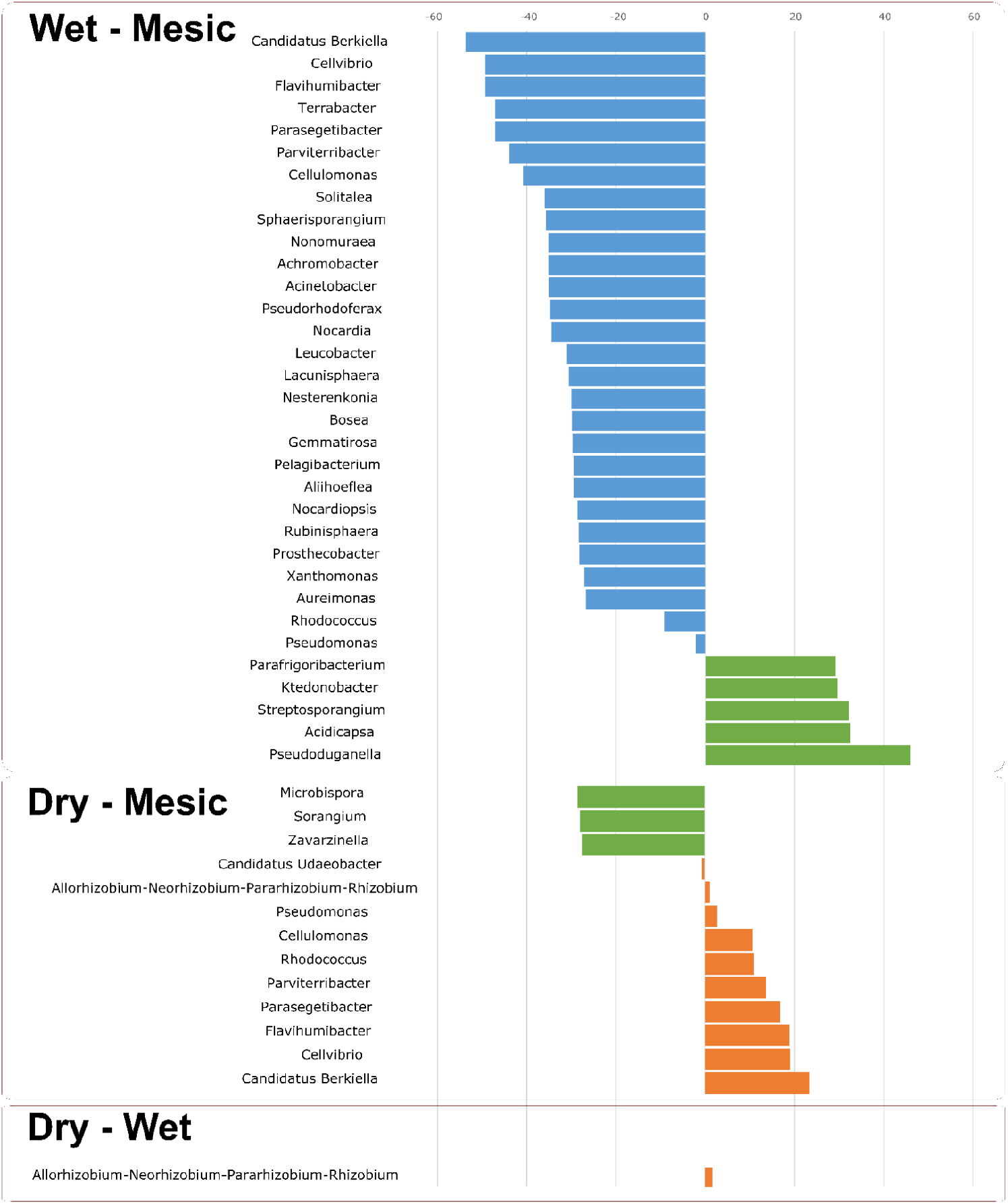
DeSEQ2 analysis to reveal the bacterial genera that were significantly different in relative abundance between the dry, mesic, and wet ecotypes (p < 0.05). Pairwise comparisons were performed between wet-mesic, dry-mesic and dry-wet ecotypes.

### No differences in ecotypic fungi diversity and structure

Unlike the observed differences in rhizobiome bacterial diversity among ecotypes, we did not notice the same pattern in the ecotypic fungi diversity and structure. We observed that the ecotypic fungal communities structure were not significantly different from each other at the phylum level (PERMANOVA, Pseudo-F=2.0827, p (perm): 0.071, p (MC): 0.08). Pairwise Kruskal-Wallis test also indicated no significant differences in the rhizosphere fungal community composition between the ecotypes (dry v wet: p (MC): 0.190, dry v mesic: p (MC): 0.117, wet v mesic: p (MC): 0.072). The top eight fungal phyla present in all three ecotypes were Ascomycota, Basidiomycota, Mortierellomycota, Mucoromycota, Glomeromycota, Chytridiomycota, Kickxellomycota, and Rozellomycota (Figure 5, Supplementary Figure S2). Besides that Ascomycota and Basidiomycota had the highest relative abundance, post-hoc SIMPER analyses at the phylum level also indicated that Ascomycota and Basidiomycota were the top phyla contributing to the similarities in all the three ecotypes (Figure 5, Supplementary Table S3). Consistent with the results shown in our study, Ascomycota dominate the rhizosphere fungal communities (Egidi et al. 2019; Praeg and Illmer 2020; Qin et al. 2017). Basidiomycota has been reported to be isolated from soil as well, and is well-known for rhizobiome dwellers (Thorn et al. 1996; Yang et al. 2017). At the genus level, *Phallus* and *Cladosporium* (SIMPER analysis) were among the top genera contributing to the similarities between the ecotypes (Supplementary Table S3).

**Figure 5:**
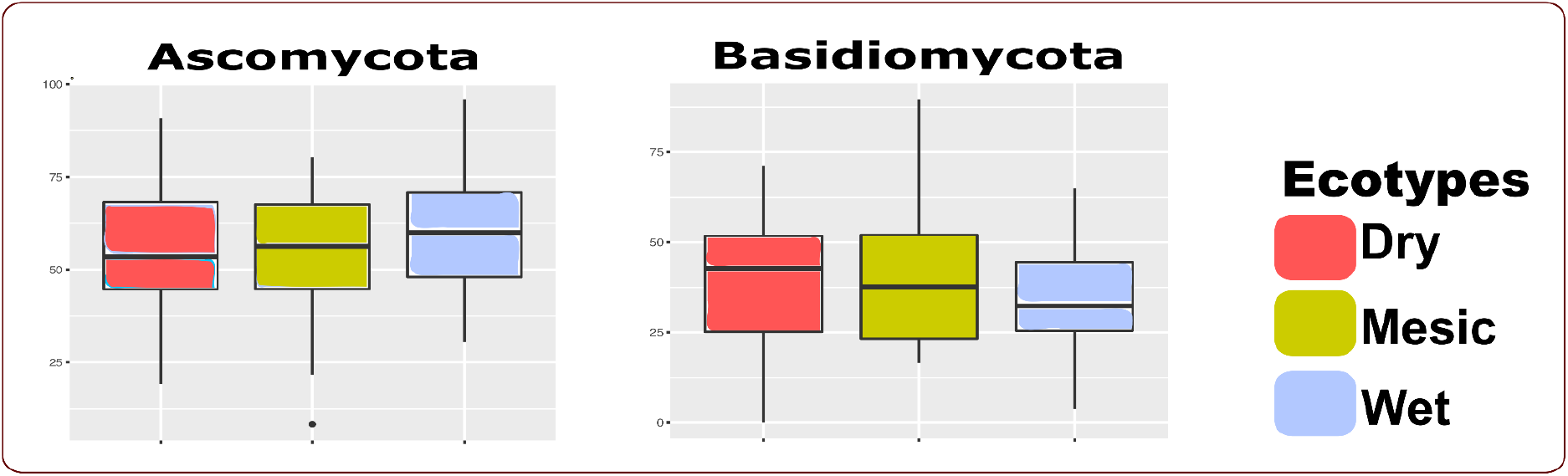
The relative abundance of Ascomycota and Basidiomycota in the dry, mesic and wet ecotypes. Ascomycota and Basidiomycota are also the most abundant phyla in all the ecotypes.

### Ecotype differences in bacterial diversity evident under arid conditions

The distribution of microorganisms in an environment is often expressed in the famous tenet “everything is everywhere but the environment selects’’ (Baas Becking, 1934). Our ten-year-long reciprocal common garden study provided an excellent opportunity to gain insights into whether and how specifically adapted ecotypic microorganisms will thrive and proliferate under arid conditions. The most ideal situation to understand these differences in bacterial diversity would include the ability to compare and contrast this study with the rhizobiome profiles before planting the *A. gerardii* ecotypes in Colby. After ten years of planting and maintaining under drought stress conditions, the environmental pressure would alter the rhizobiome structure, especially when the hosts or ecotypes had limited or no influences (Hueso, Hernández, and García 2011; Alster et al. 2013; Bachar et al. 2010; Fierer, Schimel, and Holden 2003). On the contrary, we observed that there were differences in bacterial community composition in perennial grassland ecotypes under drought stress, although there were no effects on the fungal compositions. Even without the before planting rhizobiome profiles, the bacterial community composition between ecotypes did in fact differ in the arid environment, suggesting that the host-mediated adaptation persisted even under arid conditions. The earlier report associated with *A. gerardii* has indicated that the ecotypic variation was observable in several aspects of growth in the plant. The ecotypic variation not only impacted aboveground features, but also had a prominent effect on the belowground ecosystem processes mediated by microbial communities (Mendola et al. 2015). We observed the higher relative abundance of *Rhizobium* in the dry ecotypic rhizosphere, suggesting that the *Rhizobium* might be associated with the observed higher leaf nitrogen concentrations and chlorophyll absorbance (Caudle et al.2014, Galliart et al. 2020). The plant host can implement diverse mechanisms by secreting root exudates, employing defense strategies and structural modifications to recruit a specific and optimized microbiome (Pascale et al. 2019). We demonstrated in our study that the individual plant genotype might influence the bacterial rhizobiome, and other reports showed that these traits could also be heritable (Schweitzer et al. 2008; Peiffer et al. 2013; Walters et al. 2018; Gray et al. 2014). There is a fine margin between the influence by the ecotype host and the environment on the microbiome communities. In our present work, we were able to determine that after ten years of growing under conditions outside the normal ecotypic environment, the host ecotype might be able to “overcome” environmental pressure to a certain extent and regulate the bacterial community composition. Previous studies have indicated that drought stress can escalate the relative abundance of fungal populations while decreasing the relative abundance of bacterial communities in the same poplar plantation (Sun et al. 2020). Also, it is known that bacterial networks in soil are less stable than the fungal networks (de Vries et al. 2018). Putting all these together, our study suggested that the ecotypic fungal populations might be more capable at adapting, and could have diverse resistance mechanisms towards drought conditions, resulting in the fungal community composition and structure remaining unchanged.

In this study, we explored the concept of microbial “generalists” and “specialists”. Generalist microbial populations are able to adapt to diverse habitats, while microbial populations are referred to as the specialists when those can only adapt to specific habitats (Sriswasdi, Yang, and Iwasaki 2017). Insights into the compositional analysis of the ecotypic rhizobiome suggested that the bacterial population from the dry and wet ecotypes might be specialists and are less adaptive to the induced environmental stress. We further hypothesized that the bacterial populations from the mesic ecotypes belonged to the generalist group, which strived to adapt to the arid environment. There are previous reports which acknowledge the contribution of the generalists and specialists impacting the dynamics of the structures of different microbial communities (Pandit, Kolasa, and Cottenie 2009; Székely and Langenheder 2014). We observed that the differences in bacterial community structures were more prominent between dry and mesic as well as wet and mesic, as compared to dry and wet ecotypes. Putting this all together, our study suggested that dry and wet ecotypic microbial communities might show more influences from the host as compared to that of the mesic ecotypes. From this study, we surmised that *A. gerardii* ecotypes had unique bacterial assemblage contributing to the rhizobiome. However, the classification of whether the bacterial populations were generalists or specialists might have an influence on the resultant ecotypic rhizobiome due to host and environmental interaction.

To conclude, our study provided the knowledge that will help tackle the challenges faced by grassland restorative efforts by providing insights into the impact of environmental stress on plant host-associated microbiomes. We showed that bacterial populations were influenced by the respective ecotypes, while the fungal populations were not significantly different between the ecotypes. This study also suggested the existence of host-mediated bacterial community adaptation for *A.gerardii’s* rhizosphere, and a possible trade-off between the specialist and generalist bacterial communities in specific environments, that might ultimately benefit the plant host.

## Acknowledgments

The study is based upon the work supported by the National Science Foundation EPSCoR Award No. OIA-1656006 and matching support from the State of Kansas Board of Regents. This study was supported by the United States Department of Agriculture, National Institute of Food and Agriculture (USDA NIFA), under the Award Number: 2020-67019-31803. We are thankful to Dr. Alina Akhunova of Kansas State Integrated Genomics Facility for the help with ITS and 16S amplicon sequencing.

## Supplementary Files

Supplementary Table S1: Raw sequence analysis by QIIME 2 Version 2019.7. The number of counts of bacteria and fungi initially obtained, and the counts that were considered after primer trimming and DADA2 quality control per sample.

Supplementary Table S2: Bacterial and fungal identifications along with the number of counts per sample.

Supplementary Table S3: Post-hoc SIMPER analyses indicate the bacterial and fungal phyla that contributed to major similarities among the dry, mesic, and wet ecotypes.

Supplementary Table S4: Genera that had differential relative abundance between dry, mesic and wet ecotypes.

Supplementary Figure S1: The top bacterial and archaeal taxa present in dry, mesic and wet ecotypes.

Supplementary Figure S2: The top fungal taxa present in dry, mesic and wet ecotypes.

