## Supplementary figures and images for "Bacterial but not fungal rhizosphere communities differ among perennial grass ecotypes under abiotic environmental stress"

### Supplementary Figure S1

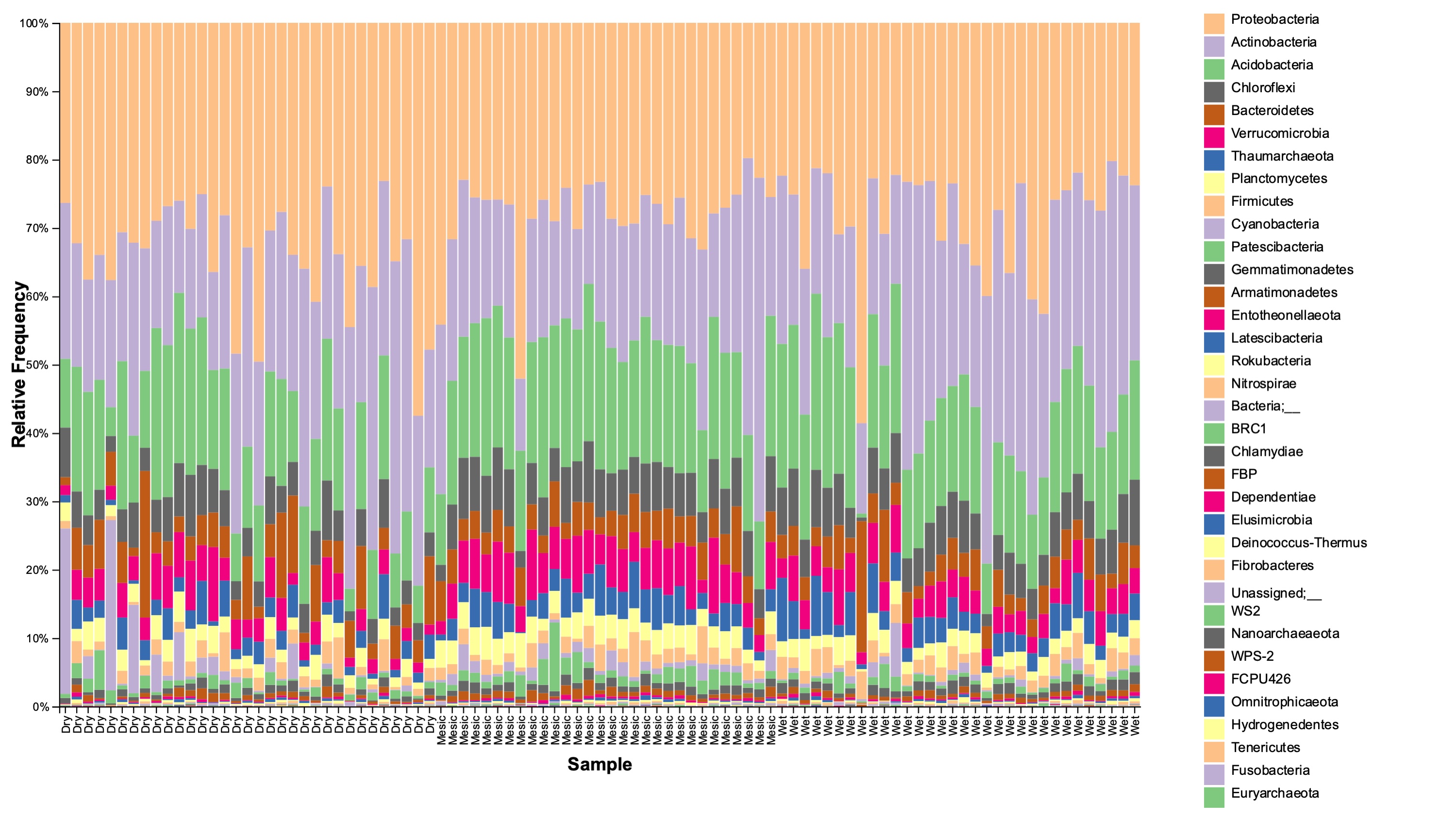

### Supplementary Figure S2

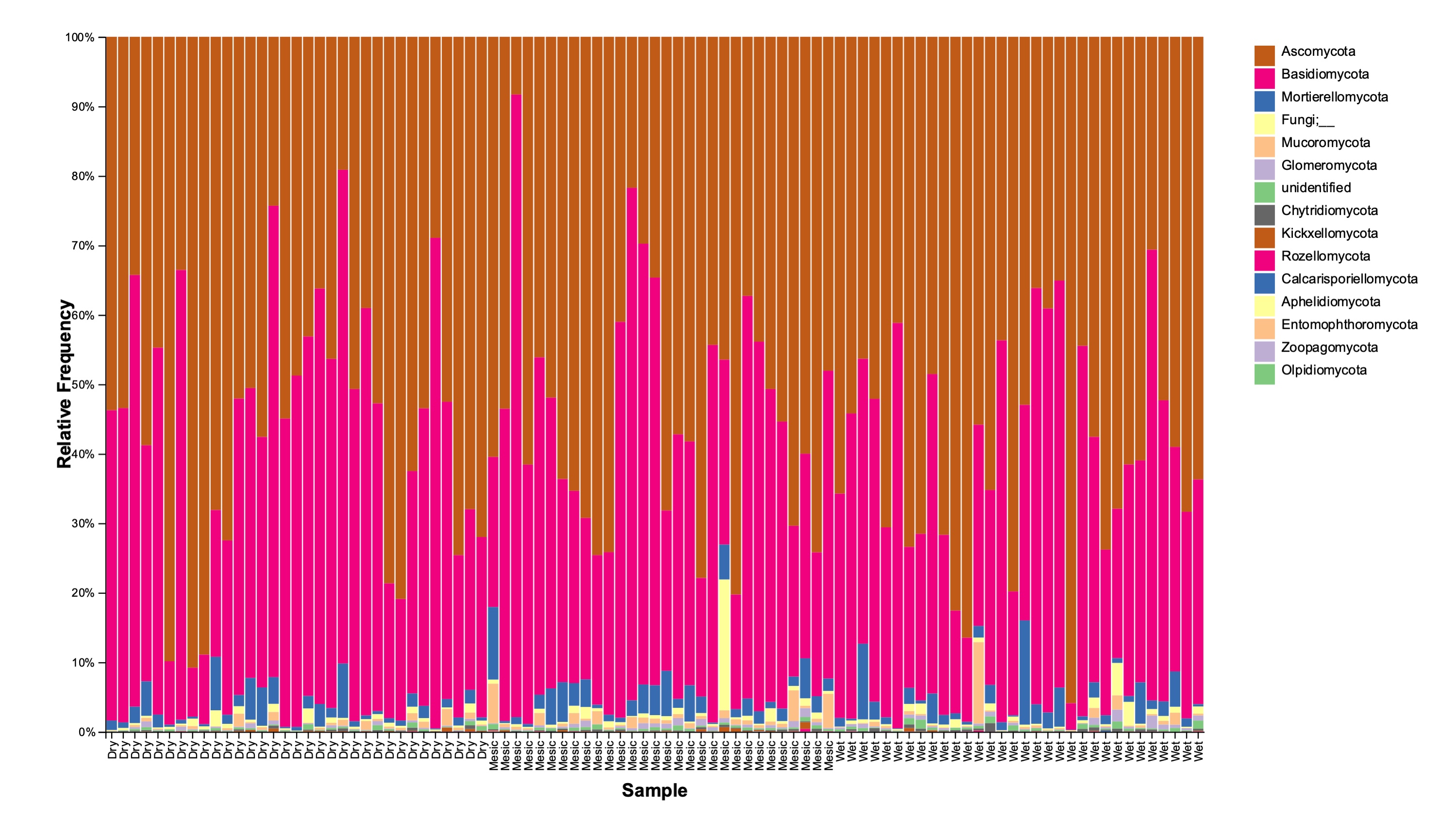
